# A novel prion strain is responsible for the first case of chronic wasting disease in Finnish moose

**DOI:** 10.1101/2022.07.11.499602

**Authors:** Julianna L. Sun, Jifeng Bian, Sehun Kim, Jenna Crowell, Bailey K. Webster, Emma K. Raisley, Sirkka-Liisa Korpenfelt, Sylvie L. Benestad, Glenn C. Telling

## Abstract

Concern is mounting over the global emergence, expanding host range, and uncertain zoonotic potential of chronic wasting disease (CWD), a fatal, infectious disease of cervids caused by prions. Our previous studies using genetically modified CWD-susceptible mice showed that Norwegian and North American CWD are caused by different prion strains. Here we investigated the properties of prions causing the first case of Finnish moose CWD. While Finnish and Norwegian moose CWD prions share characteristics that distinguish them from North American CWD including the inability to replicate in lymphoid tissues, common responses to variations at residue 226 of host prion protein, and overlapping central nervous system profiles, they also exhibit pronounced conformational variation which is consistent with strain differences between Finnish and Norwegian moose CWD. Our findings support the existence of a surprisingly diverse portfolio of emergent CWD strains in Nordic countries that is etiologically distinct from North American CWD.

**Summary line:** Prion strain properties from the first case of chronic wasting disease in a Finnish moose are similar but not identical to Norwegian cases, supporting a growing population of strains in Nordic countries.

## Introduction

Prions are infectious proteins which cause fatal, incurable neurodegenerative diseases of humans and animals that include Creutzfeldt Jakob disease (CJD); sheep scrapie; bovine spongiform encephalopathy (BSE); and chronic wasting disease (CWD) of deer, elk, moose and other cervids. The extraordinary biology and transmissibility of these disorders stems from the protean conformational properties of the prion protein (PrP). Whereas the secondary structure of normal, host-encoded cellular PrP (PrP^C^) is predominantly a-helical, during disease, PrP^Sc^, its relatively under-glycosylated infectious counterpart, assembles into amyloid fibrils with parallel in□register intermolecular b-sheets (1–4). The replicative properties of prions stem from the capacity of pathogenic PrP^Sc^ to template its infective conformation on its benign PrP^C^ counterpart in a cyclical process resulting in exponential accumulation of prion infectivity (5).

Inoculation of diseased materials into the same species typically reproduces disease with prolonged, clinically silent incubation periods. Although they lack informational nucleic acids, prions exhibit heritable strain properties which influence disease outcomes. Strain properties are defined by outcome measures of experimental transmissions in susceptible hosts and include variable times between infectious challenge and disease onset (incubation period); differences in the neurological signs defining the clinical phase; distinctive patterns of neuronal targeting and central nervous system (CNS) deposition of PrP^Sc^; and the ability to replicate in non-CNS tissues including the lymphoreticular system (LRS) and musculature (6). While prion transmission may also occur between species, this process is generally less effective than intraspecies transmission. The efficiency of interspecies transmission is influenced both by the extent of PrP sequence homology between species and by prion strain properties (7–9). Multiple lines of evidence contend that heritable strain properties are enciphered by distinct PrP^Sc^ conformations which are faithfully propagated during prion replication (10, 11).

Novel prion diseases and strain variants continue to arise in increasing numbers of animal species, and the food chain transmission of BSE to humans resulting in a variant of CJD illustrates the unpredictable potential of animal prion diseases for zoonotic transmission (9). The burgeoning epidemic of CWD is of particular concern because of its unparalleled contagious transmission and expanding host range in wild as well as captive cervids, and the identification of increasing numbers of CWD strains with increasingly uncertain zoonotic potential (12). Following its initial description in captive deer (13), the ensuing spillover to free-ranging deer and elk was originally confined to an endemic region in Colorado and Wyoming. Subsequent uncontrolled transmission has resulted in growing numbers of CWD-affected cervids in at least 28 American states and three Canadian provinces. CWD is also now endemic in the Republic of Korea where it infected new cervid species following inadvertent importation of subclinically diseased animals from North America (NA) during the 1990’s (14–16). Following the diagnosis of CWD in a free-ranging Norwegian reindeer in 2016 (17), additional cases were subsequently diagnosed in growing numbers of moose, red deer, and reindeer in Norway, Sweden, and Finland (18).

While experimental limitations of natural host species hampered initial characterizations of CWD, the development and application of susceptible mice expressing cervid PrP^C^ (CerPrP^C^) provided insights into multiple aspects of pathogenesis, including the mechanism by which naturally-occurring *PRNP* coding sequence variations impact the selection and propagation of CWD strains (12). Cervid *PRNP* coding sequences vary at codon 226: whereas deer, reindeer, and moose encode glutamine (Q) (CerPrP-Q226), elk or red deer may encode glutamate (E) at this position (CerPrP-E226). In order to precisely assess the effects of this primary structural difference we created CWD-susceptible gene targeted (Gt) mice in which the murine PrP coding sequence was targeted and precisely replaced with that of CerPrP-Q226 or CerPrP-E226, referred to as GtQ and GtE mice (19). Since GtE and GtQ express accurately controlled, equivalent levels of CerPrP^C^ and are syngeneic except at codon 226, we were able to ascribe distinct CWD outcomes in GtQ and GtE mice to amino acid variation at this position (19). This approach not only ascribed a key role for this primary structural difference in CWD strain selection, but also underscored the utility of Gt mice for accurately defining the strain properties of emergent CWD prions (20). Moreover, in contrast to previously produced transgenic (Tg) mice, Gt mice also recapitulated the lymphotropic properties of CWD strains (19, 20). Using Gt mice we showed that emergent Norwegian and established NA forms of CWD are caused by unrelated prion strains that respond differently to variation at residue 226 and have different lymphotropic properties (20). The goals of our current studies were to assess the strain properties of prions causing a newly emergent form of CWD in moose from Finland, and to compare them with those of NA and Norwegian CWD.

## Results

We intracerebrally inoculated GtE and GtQ mice and their overexpressing Tg counterparts (TgE and TgQ) (21, 22) with homogenates of frozen CNS and LRS materials from the first diagnosed case of CWD in moose from Finland, referred to here as M-F1. CNS homogenates produced disease in ~ 90 % of GtQ mice after ~ 250 d and 75 % of TgQ mice after ~ 230 d (Fig. 1A, Table 1). In contrast, all inoculated GtE mice remained free of disease for > 550 d, and only a single TgE mouse developed clinical signs after ~ 400 d (Fig. 1A and Table 1). Western blotting of proteinase K (PK)-resistant CerPrP^Sc^ in mouse brains confirmed the diagnoses of prion disease (Fig. 2B). In contrast, LRS homogenates failed to produce disease in mice expressing either CerPrP^C^-Q226 or CerPrP^C^-E226 after elapsed times approaching 600 d (Fig. 1B, Table 1). We deduce that CWD prions failed to propagate in the LRS of Finnish moose M-F1. We also conclude that propagation of CNS-derived M-F1 CWD prions is favored in mice expressing CerPrP^C^-Q226 and relatively restricted in mice expressing CerPrP^C^-E226.

**Fig. 1:**
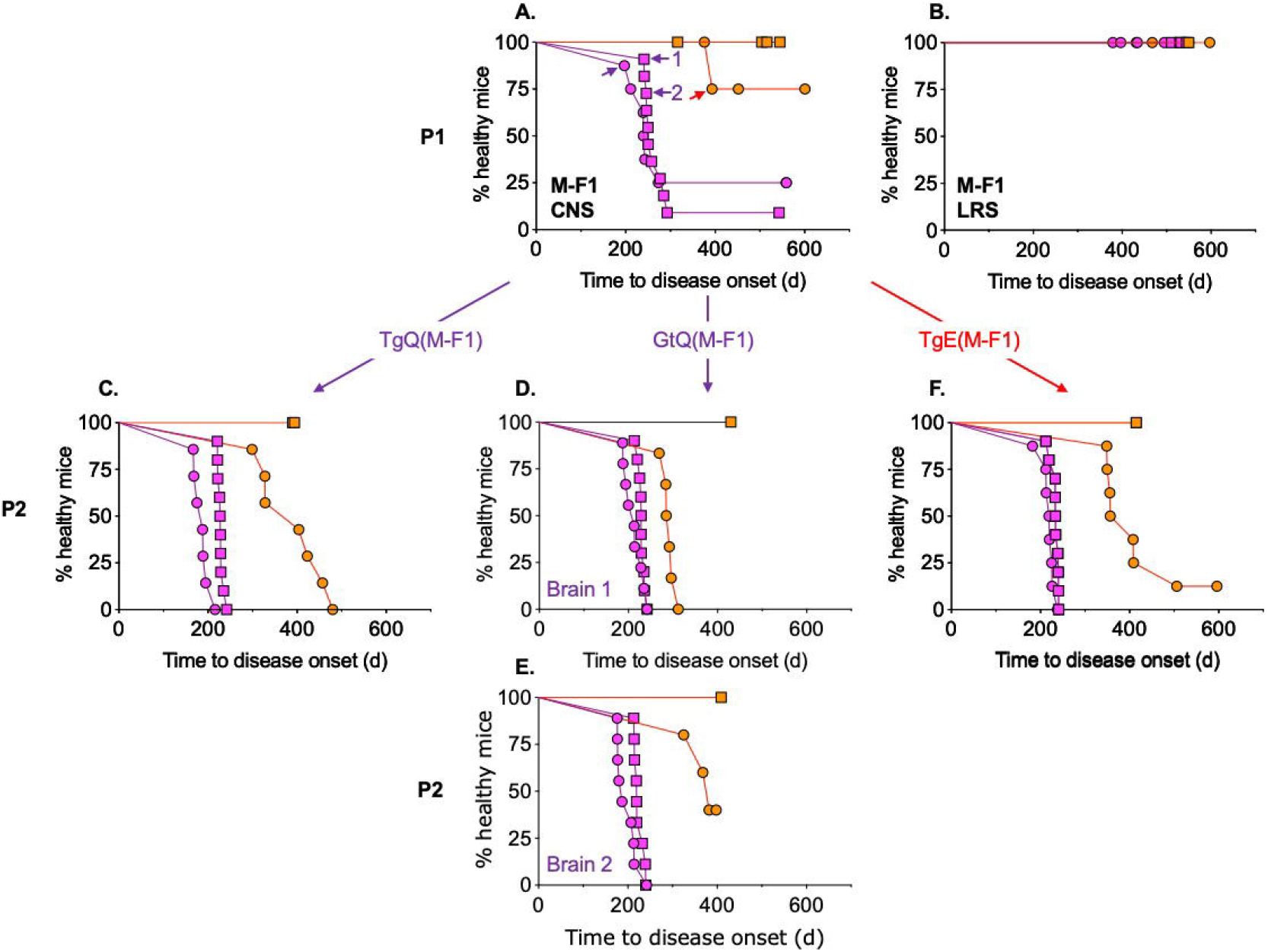
Transmission properties of CWD prions from the fist diagnosed case of CWD in Finland. Survival curves of TgE (orange circles), TgQ (magenta circles), GtE (orange squares), and GtQ mice (magenta squares) following inoculation with homogenates of **A.** frozen CNS or **B.** LRS tissues from Finnish moose M-F1. In **A.**, the TgQ mouse and the two GtQ mice (1 and 2) from which brains were used for serial transmission are indicated by arrows. **C.**, serial passage of M-F1 prions from TgQ. **D.** and **E.**, serial passage of M-F1 prions from two GtQ mice. **F.**, serial passage of M-F1 prions from TgE.

**Fig. 2:**
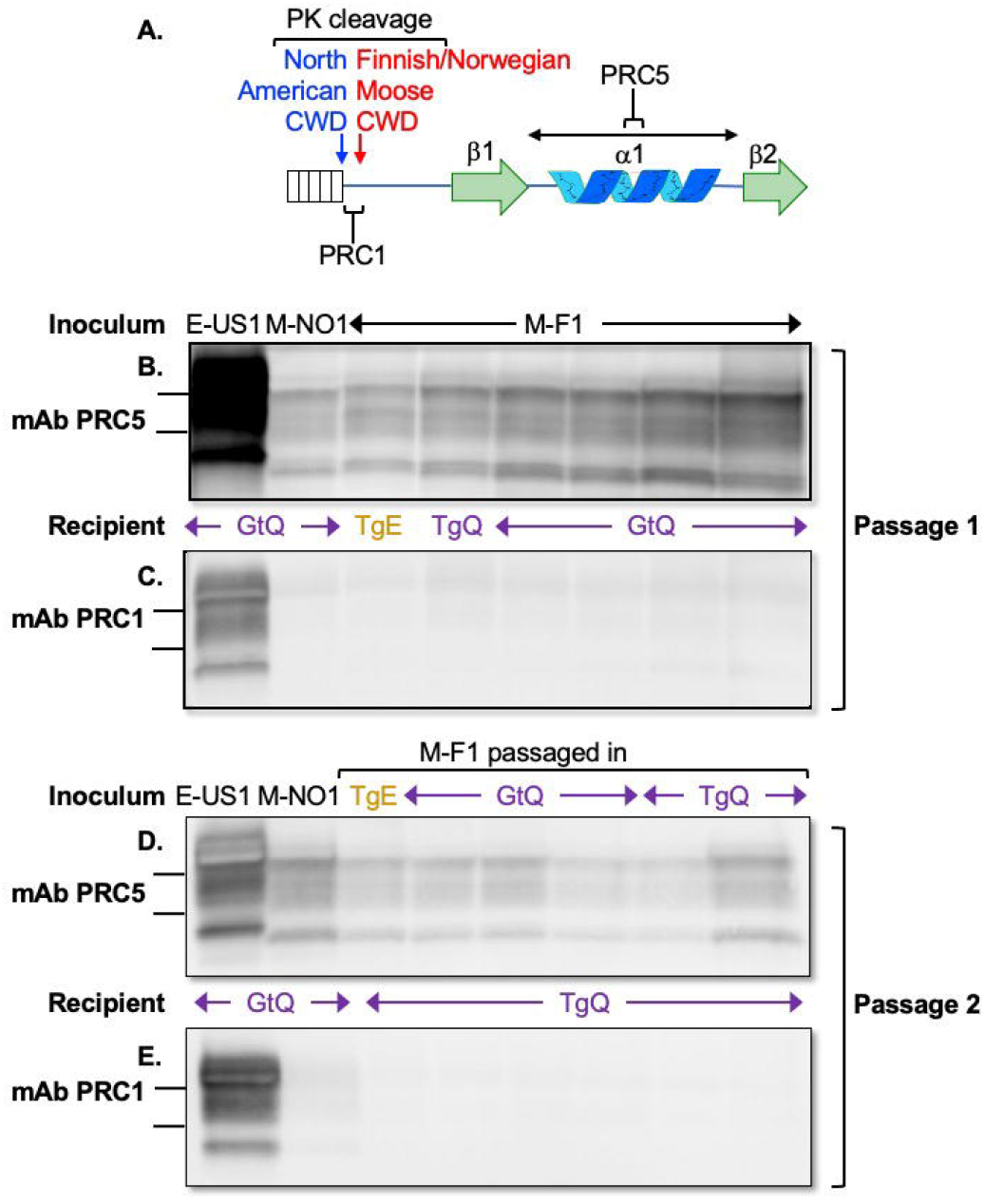
Western blot comparisons of CerPrP^Sc^ in the CNS of mice infected with Nordic and North American CWD. **A.** Location of PrP epitopes for mAbs PRC5 and PRC1 and inferred PK cleavage sites for NA and Finnish moose CWD. **B.** and **C.**, western blots of CNS homogenates from GtQ, TgQ and TgE mice after primary transmissions of elk CWD from USA (E-US1), Norwegian moose CWD (M-NO1), or Finnish moose CWD (M-F1) probed with mAb PRC5 (**B.**) or PRC1 (**C.**). **D**. and **E.**, western blots of CNS homogenates of GtQ and TgQ mice infected with M-F1 passaged through TgQ, GtQ or TgQ, as indicated, and NA E-US1 and Norwegian M-NO1 passaged to GtQ mice probed with mAb PRC5 (**D.**) or PRC1 (**E.**).

**Table 1:** Susceptibility of transgenic and gene-targeted mice intracerebral challenges with Finnish and Norwegian moose CWD prions.

| **A. Finnish moose CWD** |  |  |  |  |  |  |
| --- | --- | --- | --- | --- | --- | --- |
| Inoculum | Source tissue | Passage | Mean incubation times (days), ± SEM (n/n<sub>0</sub>) | | | |
|  |  |  | TgE | TgQ | GtE | GtQ |
| M-F1 | CNS | p1 | 393 (1/5) | 234 ± 27 (6/8) | >545 (0/9) | 259 ± 19 (10/11) |
| | CNS (TgQ) | p2 | 388±27 (7/7) | 182 ± 7 (8/8) | >403 (0/6) | 228 ± 3 (10/10) |
| | CNS (GtQ 1) | p2 | 290±6 (6/6) | 205 ± 9 (10/10) | >424 (0/8) | 228 ± 3 (10/10) |
| | CNS (GtQ 2) | p2 | 374±13 (5/5) | 197±8 (9/9) | >403 (0/5) | 223±4 (9/9) |
| | CNS (TgE) | p2 | 390 ± 22 (7/8) | 217 ± 6 (8/8) | >424 (0/8) | 232 ± 3 (10/10) |
|  | LRS | p1 | >597 (0/9) | >550 (0/9) | >550 (0/6) | >545 (0/10) |

| **B. Norwegian moose CWD** |  |  |  |  |  |  |
| --- | --- | --- | --- | --- | --- | --- |
| M-NO1 | CNS | p1 | 570 ± 39 (4/5) | 297 ± 43 (9/9) | >553 (0/7) | 440 ± 13 (13/13) |
| | CNS (TgQ) | p2 | 337 ± 51 (6/8) | 181 ± 27 (10/10) | 540 ± 29 (2/5) | 271 ± 30 (8/8) |
| | CNS (GtQ) | p2 | | | >524 (0/8) | 281 ± 24 (5/5) |

Time to disease onset (incubation time) is expressed as the mean, in days, at which inoculated mice first developed ultimately progressive signs of neurological disease ± standard error of the mean (SEM). n/no, number of diseased mice/number of inoculated mice.

Iterative passages of M-F1 prions from diseased TgQ mice elicited disease in GtQ mice after ~ 230 d and in TgQ mice after ~ 180 d (Fig. 1C, Table 1). Serial propagation of M-F1 prions from the brains of two diseased GtQ mice also caused disease in all inoculated GtQ and TgQ mice after ~ 200 to 230 d (Fig. 1D and E, Table 1). CerPrP^C^-E226-expressing mice were less responsive. Currently, no disease has been registered in GtE mice inoculated with TgQ-passaged M-F1 after > 410 d, which is ~ 180 d beyond the time to disease in GtQ mice (Fig. 1C; Table 1), and disease in TgE mice occurred at ~ 380 d, ~ 180 d after the mean time to disease in TgQ mice (Fig. 1C; Table 1). GtE and TgE mice were similarly poorly responsive to M-F1 from the brains of two diseased GtQ mice (Fig. 1D, E; Table 1). We also serially passaged prions from the CNS of the single TgE mouse which developed disease after primary challenge with M-F1 (Fig. 1A; Fig. 2B; Table 1). Whereas TgE-passaged M-F1 prions produced disease in GtQ mice after ~ 230 d, they have currently failed to produce disease in GtE mice after > 430 d. Similarly, while TgE-passaged M-F1 prions produced disease in TgQ mice within ~ 170 d, only 90 % of inoculated TgE mice developed disease after ~ 390 d, (Fig. 1F; Table 1). Collectively, these serial passing results support our conclusion that CerPrP^C^-Q226 provides an optimized template for conversion of M-F1 CWD prions while expression of CerPrP^C^-E226 restricts their transmission.

Comparison of the transmission profiles of Finnish and Norwegian moose CWD (20) revealed that primary and secondary transmissions of both M-F1 and Norwegian M-NO1 prions were more effective in mice expressing CerPrP^C^-Q226 than in CerPrP^C^-E226 expressing counterparts. However, mean incubation times of serially passaged M-F1 and M-NO1 to GtQ mice differed by ~ 40 to 50 d (*P* > 0.001 – 0.0001) (Table 1). These discrepant incubation times suggest that, despite their broadly similar responses to variation at residue 226, the strain properties of these Finnish and Norwegian CWD prions are not completely concordant.

Using western blotting, we compared the properties of CerPrP^Sc^ constituting M-F1-in infected Tg and Gt mouse brains with those of NA and M-NO1 CWD. Using monoclonal antibody (mAb) PRC5 which recognizes an epitope in the structured globular domain of PrP (23) (Fig. 2A), we found that the electrophoretic migration of M-F1 was more rapid than NA CWD, and equivalent to the previously-reported faster migrating M-NO1 (20) (Fig. 2B). Accumulation of M-F1 proteinase resistant CerPrP^Sc^ was reduced compared to NA CWD, and these lower levels were similar those previously-reported for M-NO1 (20) (Fig. 2B). Also in accordance with M-NO1 (20), M-F1 was relatively refractory to detection by PRC1, while NA CWD was reactive with this mAb (Fig. 2C). Since the PRC1 epitope is immediately distal to the site of PK cleavage in NA CerPrP^Sc^ (23) we conclude that the faster migration and resistance of M-F1 and M-NO1 to PRC1 detection results from epitope loss due to PK cleavage downstream from the site in NA CWD (Fig. 2A). Serial transmission of M-F1, M-NO1 and NA CWD from TgE, GtQ, and TgQ produced similar immunoblotting profiles and differential reactivity with mAb PRC1 (Fig 2D, E).

We assessed the responses of M-F1, M-NO1 and NA CWD prions to denaturation with increasing concentrations of guanidine hydrochloride (GdnHCl) (Fig. 3). This measure of PrP^Sc^ stability is associated with conformational variation among prion strains (24). Stability profiles and concentrations of guanidine hydrochloride (GdnHCl) producing half-maximal denaturation (GdnHCl_1/2_) indicated that the stability of M-F1 in GtQ mice was substantially lower than M-NO1 (*P* > 0.0001), and equivalent to NA moose CWD (Fig. 3A). Similar conformational stabilities of M-F1 were observed in TgQ mice, and following infection of TgQ mice with M-F1 previously passaged in TgE or TgQ mice. Denaturation curves of these prions were overlapping and GdnHCl_1/2_ values were comparable in the range of 2.59 to 2.95 (Fig. 3B). We conclude that conformational properties of M-F1 prions are distinct from those of M-NO1 prions, and that the conformation of M-F1 prions remains unaltered following transmissions to mice expressing CerPrP^C^-E226 and CerPrP^C^-Q226.

**Fig. 3:**
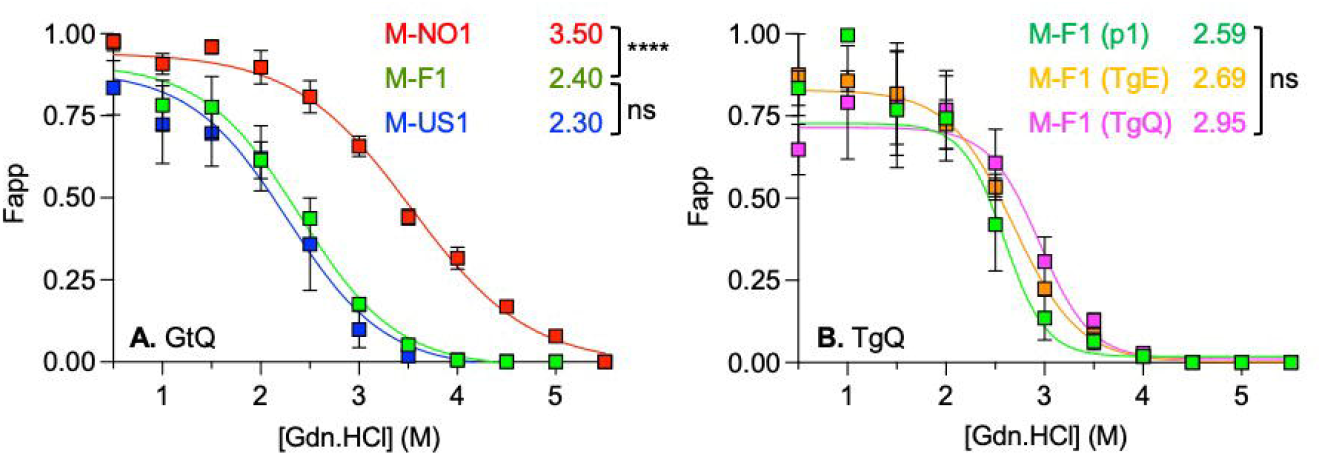
Responses of Nordic and North American moose CWD prions to increasing concentrations of guanidine hydrochloride. **A.** Responses of PK-resistant CerPrP^Sc^ constituting moose CWD prions to denaturation with increasing concentrations of GdnHCl. Green symbols, Finnish moose M-F1; red symbols, Norwegian moose M-NO1; blue symbols, US moose M-US1. **B.** Responses of M-F1 CerPrP^Sc^ from the CNS of TgQ mice to denaturation with increasing concentrations of GdnHCl following passage of M-F1 through TgQ and TgE mice. Green symbols, M-F1 in TgQ mice; orange symbols M-F1 passaged through TgE mice; magenta symbols, M-F1 passaged through TgQ mice. PK-resistant PrP^Sc^ was quantified by densitometry of immunoprobed dot blots and plotted against GdnHCl concentration. Sigmoidal dose-response curves were plotted using a four-parameter algorithm and nonlinear least-square fit. F_app_, fraction of apparent PrP^Sc^ = (maximum signal-individual signal)/(maximum signal-minimum signal). Error bars, ± standard error of the mean of data from analyses of three animals per group. GdnHCl_1/2_ values (M) for each infection are reported on the right-hand side of each graph. Significance calculated by pairwise analysis of GdnHCl_1/2_ values from best fit curves.

Histoblotting of coronal brain sections (25) showed that the deposition of CerPrP^Sc^ in M-F1 infected GtQ mice was widespread, diffuse, and symmetrically distributed across both brain hemispheres (Fig. 4A). This distribution pattern was recapitulated in M-F1-infected TgQ mice (Fig. 4B), and after iterative passage to additional GtQ mice (Fig. 4C). The pattern of M-F1 distribution was indistinguishable from that of M-NO1 in GtQ mice (Fig. 4D). These M-F1 and M-NO1 patterns differed from the disorganized, asymmetrically distributed amalgamations of NA CWD in GtQ mice (Fig. 4E) (20).

**Fig. 4:**
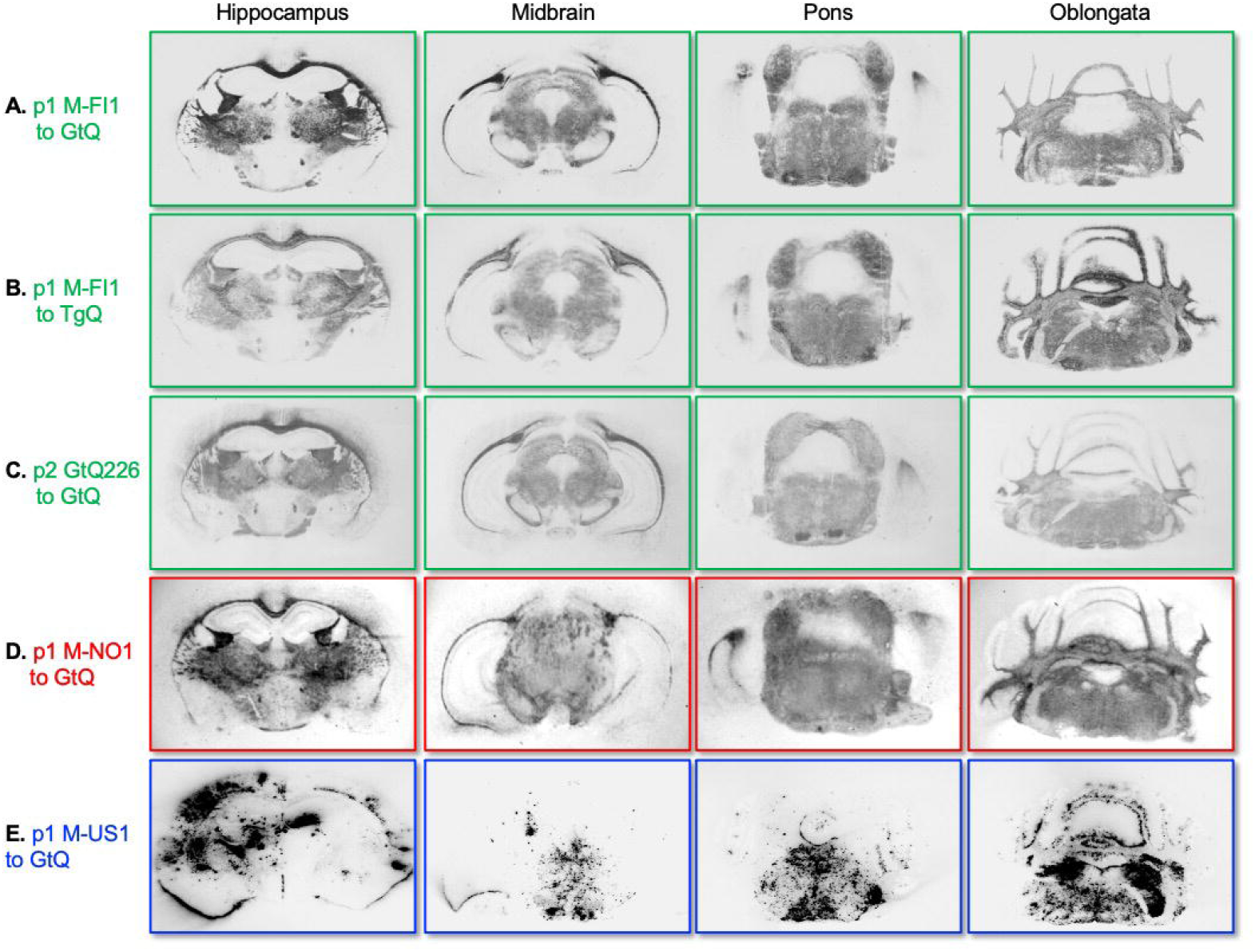
Global CNS distribution of CerPrP^Sc^ in Tg and Gt mice infected with Nordic and North American CWD. Cryostat coronal brain sections were taken at the level of the hippocampus, midbrain, pons, and oblongata and transferred to slides and then to nitrocellulose. Sections were PK treated, and immunoprobed with mAb PRC5 after denturation. **A.B.** M-F1 passaged in GtQ and TgQ mice respectively. **C.** GtQ-passaged M-F1 serially passaged in GtQ mice. **D.** M-NO1 passaged in GtQ mice. **E.** M-US1 passaged in GtQ mice. PK treated histoblots were probed with mAb PRC5.

Microscopic analysis of CNS sections from infected mice revealed the presence of spongiform degeneration (Fig. 5). These sections were immunohistochemically treated to reveal disease-associated CerPrP after reaction with anti-PrP antibodies. Small punctate accumulations set against a background of diffuse staining were visible at high magnification of CNS sections from M-F1-infected GtQ mice (Fig. 5A). M-F1-infected TgE mice revealed a similar pattern of accumulation (Fig. 5B). Only small punctate accumulations were detected in M-NO1-infected GtQ mice (Fig. 5C). In contrast, and consistent with previous findings (19, 20), GtQ mice infected with NA CWD contained intensely staining, amorphous aggregates of CerPrP-Q226 (Fig. 5D).

**Fig. 5:**
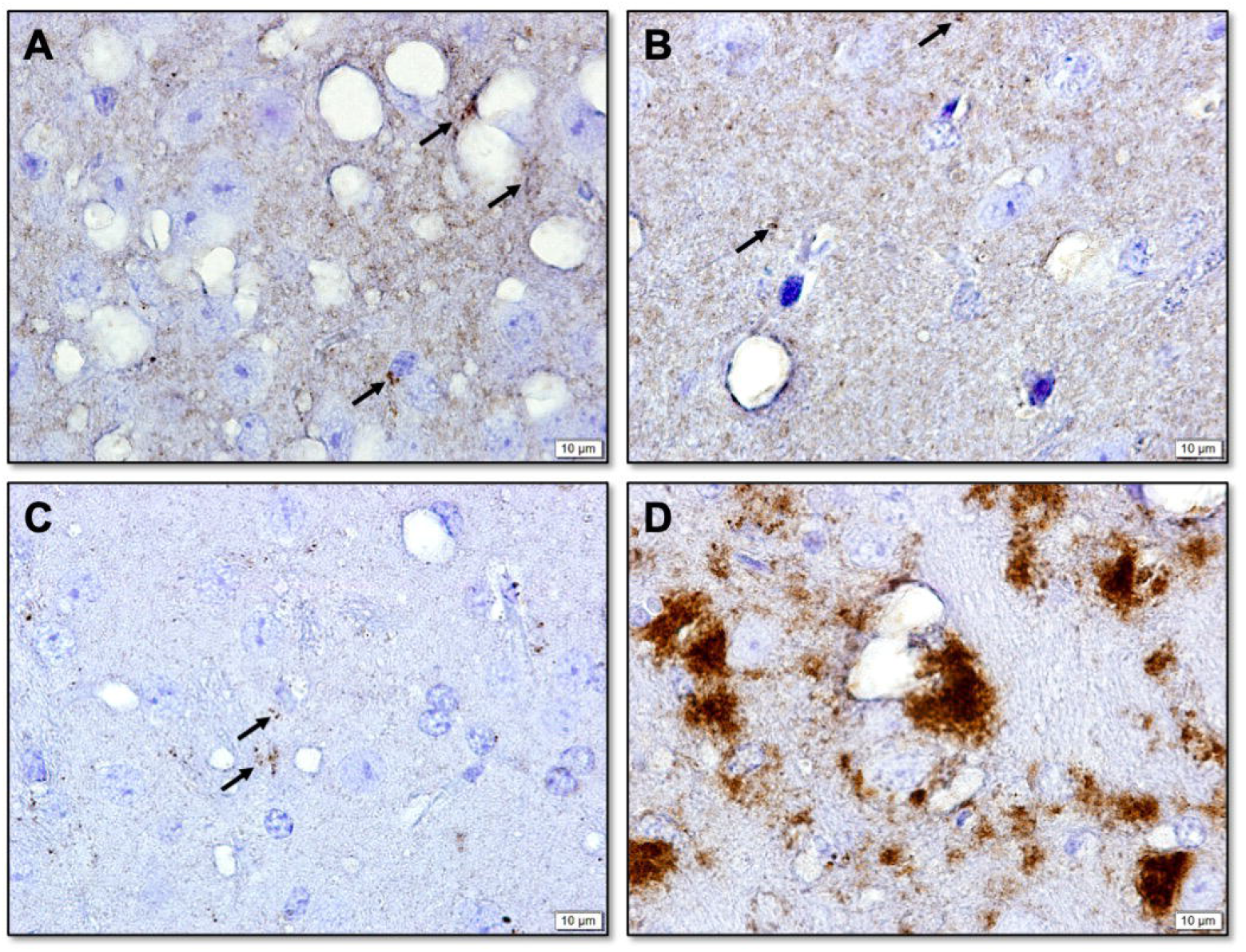
Immunohistochemical analysis of disease-associated PrP in the CNS of Tg and Gt mice infected with Nordic and North American CWD. **A.** GtQ mice infected with M-F1; **B.** TgE mice infected with M-F1; **C.** GtQ mice infected with M-NO1; **D.** GtQ mice infected with M-US1. Arrows in **A.**, **B.**, and C. point to small puncta of PrP. IHC sections were stained with Fab D18. Scale bar = 10□M.

## Discussion

Our previous studies revealed broadly similar transmission properties of NA moose, deer and elk CWD prions including their responses to amino acid variation at CerPrP residue 226. While Gt and Tg mice expressing either CerPrP^C^-E226 or CerPrP^C^-Q226 were both susceptible to NA CWD, times to disease onset were consistently more rapid in Gt and Tg mice expressing CerPrP^C^-E226 than their CerPrP^C^-Q226-expressing counterparts (19, 20, 22, 26). In contrast, transmission of emergent Norwegian moose CWD produced long incubation times and incomplete attack rates in Gt and Tg mice expressing CerPrP^C^-E226, and efficient transmission was only registered in Gt and Tg mice expressing CerPrP^C^-Q226 (20). Those studies revealed at least two strain classes of moose CWD; one defined by the transmission properties of M-NO1; the other by those of M-NO2 and M-NO3. These distinct moose CWD strains were different again from those causing CWD in Norwegian reindeer (20). The different properties of all emergent Norwegian strains compared to NA CWD made it unlikely that they were causally related. Experiments performed in parallel in bank voles supported this interpretation (27). Clear differences in the strain properties of M-F1 and NA CWD reported here allow us to build on our previous conclusions derived from studies of Norwegian CWD (20, 27), and to state more broadly that the etiology of Nordic CWD is distinct from NA CWD.

Our analyses reveal that certain characteristics of M-F1 CWD prions overlap with those of Norwegian moose CWD. Like Norwegian moose CWD isolates, propagation of M-F1 was more efficient in mice expressing CerPrP^C^-Q226 than in mice expressing CerPrP^C^-E226. We take this to mean that replication of CWD strains from Norwegian and Finnish moose is favored by expression of CerPrP^C^-Q226 and restricted by CerPrP^C^-E226. Western and histoblot profiles of M-F1 and M-NO1 CerPrP^Sc^ are also indistinguishable. Our previous studies showed that, unlike NA CWD prions which are lymphotropic (19, 28), Norwegian moose CWD prions are non-lymphotropic and fail to accumulate in the spleens of infected GtQ mice (20). Our current findings that Gt or Tg mice are resistant to disease when challenged with LRS tissue preparations from M-F1 indicates that prions were confined to the CNS of this Finnish moose, and therefore that M-F1 prions, like those causing CWD in Norwegian moose, are also non-lymphotropic.

Despite these broadly similar characteristics, other measures are consistent with strain variation between M-F1 and M-NO1 CWD. Differences supporting this conclusion include serial passage outcomes in GtQ mice, subtle variations in IHC profiles of disease-associated PrP, and, most strikingly, the responses of M-F1 and M-NO1 prions to GdnHCl denaturation which reveal distinct stabilities consistent with pronounced strain-dependent conformational differences. These incompletely overlapping properties of Finnish and Norwegian moose CWD prions add to an increasing body of evidence for a surprising variety of strains among Nordic cervids which stands in contrast to a relatively consistent CWD strain profile among NA deer, elk and moose. As yet, there appear to be no clear insights into the reasons for this unexpectedly diverse prevalence of Nordic CWD strains, or indeed their origins, although sporadic disease and prion transmission from other species have been considered as possibilities (20). Efficient prion dissemination to peripheral tissues of affected cervids has been cited as a reason for the contagious properties of NA CWD (28). While the non-lymphotropic properties of Finnish and Norwegian moose CWD strains therefore appear to militate against contagious propagation from infected Nordic moose, our descriptions of an increasing incidence of CWD strains with divergent strain properties in moose, reindeer, and red deer emergent in a relatively confined geographic location during a short time also presents challenges when considering an sporadic etiology. Four moose from Sweden have also shown evidence of CerPrP^Sc^ accumulation (18). Analyses of the strain properties of these additional emergent CWD cases will be of considerable interest.

Two other findings from this study are worthy of discussion and build upon our previous observations. Seminal studies indicated that related PrP^Sc^ and PrP^C^ primary structures supported optimal prion propagation, and this concept laid the foundation for the development of a wide array of genetically modified mice which were predictably susceptible to diverse human and animal prion diseases (8, 29, 30). Accordingly, since moose express CerPrP^C^-Q226, and CWD prions from this species are consequently composed of CerPrP^Sc^ with Q at residue 226 (CerPrP^Sc^-Q226), we reasoned that the relatively inefficient transmission of M-F1 in GtE and TgE mice could be overcome by acclimatization in mice expressing CerPrP^C^-E226, resulting in adapted M-F1 prions composed of CerPrP^Sc^-E226. However, our findings showing comparable transmission profiles of TgE-adapted M-F1 prions and M-F1 moose prions composed of CerPrP^Sc^-Q226 indicate that M-F1 prions favor conformational conversion of CerPrP^C^-Q226 over CerPrP^C^-E226 regardless of whether they are composed of CerPrP^Sc^-E226 or CerPrP^Sc^-Q226. The inability of M-F1 prions to efficiently adapt to a host expressing a single primary structural difference is reminiscent of our findings showing that the induction of prion disease by particular strains in hosts expressing new PrPC primary structures does not inevitably result in their complete adaptation and subsequent sustained propagation in that new host. Instead, prions that arise from this process, referred to as non-adaptive prion amplification (NAPA), retain atavistic preferences for their species of origin (31). Thus, while M-F1 prions are capable of sub-optimal propagation by NAPA using CerPrP^C^-E226 as template for replication, the resulting M-F1(E226) NAPA prions are not adapted for further propagation by CerPrP^C^-E226, but instead retain a preference for conversion of CerPrP^C^-Q226.

Of final note, although early studies in Tg mice supported an inverse relationship between prion incubation times and levels of PrP transgene expression (29), M-F1 prions produced disease in GtQ mice with similar mean incubation times to TgQ mice, despite expressing ~ five-fold lower levels of CerPrP^C^-Q226 expression. These findings are consistent with our previous comparative transmissions of CWD prions in Tg and Gt mice (20). In light of the precise and authentic control of PrP expression, and the ability of Gt mice to recapitulate additional cardinal features of CWD including strain-dependent lymphotropism, our findings support the general notion that targeted replacement of the murine PrP coding sequence in *Prnp* offers a superior platform for studying manifold aspects of prion biology, including mechanisms of strain selection and adaptation in peripheral compartments as well as the CNS, and the role of these processes in intra- and inter-species transmission of prion diseases.

## Materials and Methods

### Ethics Statement

All animal work was performed in an Association for Assessment for Accreditation of Laboratory Animal Care accredited facility in accordance with the Guide for the Care and Use of Laboratory Animals. All procedures used in this study were performed in compliance with and were approved by the Colorado State University Institutional Animal Care and Use Committee.

### CWD isolates

The Finnish moose isolate, M-F1, was found in 2018 in a free-ranging female moose identified to be sick and later found dead. We thank Marja Isomursu for CWD surveillance and sample collection in Finland. The NA moose isolate, M-US1, was a captive moose orally inoculated with a pooled preparation of deer CWD. The NA elk CWD isolate, E-US1, and the Norwegian moose CWD isolate, M-NO1, have been described previously (20). *Mouse models and incubation time assay* The development and characterization of TgQ, TgE, GtQ, and GtE mice and prion incubation time assay have been described previously (19–22). Equal numbers of male and female mice were used in inoculation studies. Clinical diagnoses of prion disease in mice were confirmed by PrP^Sc^ detection in the CNS.

### Analysis of PrP^Sc^ by western blotting

was performed as described previously, for example (19, 20). SDS-PAGE was performed using precast 12 % discontinuous Bis-Tris gels (Bio-Rad Laboratories, Hercules, CA). Proteins transferred to PVDF-FL membranes (Millipore, Billerica, MA, USA) were detected with mAb PRC5 at a dilution of 1:5000 and mAb PRC1 at a dilution of 1:2500 (23), followed by horse radish peroxidase (HRP)-conjugated anti-mouse IgG secondary antibody. Membranes were developed using ECL 2 western blot substrate (Thermo Scientific, USA).

### Conformational stability assay

was conducted as previously described (19, 20). PrP^Sc^ on membranes was detected with mAb PRC5 at a dilution of 1:5000, followed by HRP-conjugated goat anti-mouse IgG secondary antibody. Membranes were developed with ECL 2 western blot substrate and scanned with an ImageQuant LAS 4000 (GE Healthcare), and signals analyzed with ImageQuant TL 7.0 software (GE Healthcare).

### Histoblot analysis

was performed as previously described (25). PrP^Sc^ on membranes was detected using mAb PRC5 at a dilution of 1:5000 followed by alkaline phosphatase conjugated goat anti mouse IgG (Southern Biotech, 1030-04) at a dilution of 1:5000. Membranes were developed using 5-bromo-4-chloro-3-indolyl phosphate (BCIP)/nitro blue tetrazolium (NBT) (Sigma Aldrich, 11697471001. Images were captured using a Nikon Z1000 microscope.

### Immunohistochemical analyses

Slides were heated to 60 ? C for 30 min prior to xylene and graduated ethanol treatment followed by treatment with 88 % formic acid for 30 min. Antigen retrieval was performed in the 2100 Retriever (ProteoGenix, Schiltigheim, France) using citrate buffer followed by endoperoxidase quenching in 3 % hydrogen peroxide. Slides were blocked in 5 % non-fat milk for 30 min at room temperature before overnight incubation at *4□* C with primary antibody D18 at a 1:2500 dilution. Slides were exposed to biotin labelled goat Fab anti-human IgG secondary antibody (Southern Biotech, Birmingham, AL) at a 1:2500 dilution for 1 h at room temperature, then developed for 30 min at room temperature utilizing avidin-conjugated HRP with diaminobenzidine (DAB) as substrate (Vector Laboratories, Burlingame, CA). Slides were counterstained with hematoxylin, run through graduated ethanol treatment, cover slipped, and imaged at 100 x under oil immersion.

### Statistical information

Statistical analyses were performed using GraphPad Prism software (San Diego, CA). Statistical significance between survival curves of inoculated groups was assessed by comparing median times of survival of various inoculated groups using the log rank (Mantel-Cox) test. Statistical significance between denaturation curves of PrP^Sc^ was assessed by comparing log EC_50_ values.

## References

1. Riek R, Hornemann S, Wider G, Billeter M, Glockshuber R, Wüthrich K. NMR structure of the mouse prion protein domain PrP(121-231). Nature. 1996;382:180–2.

2. Kraus A, Hoyt F, Schwartz CL, Hansen B, Artikis E, Hughson AG, et al. High-resolution structure and strain comparison of infectious mammalian prions. Mol Cell. 2021 Nov 4;81(21):4540–51 e6.

3. Kang H-E, Bian J, Kane SJ, Kim S, Selwyn V, Crowell J, et al. Incomplete glycosylation during prion infection unmasks a prion protein epitope that facilitates prion detection and strain discrimination. The Journal of biological chemistry. 2020.

4. Basler K, Oesch B, Scott M, Westaway D, Wälchli M, Groth DF, et al. Scrapie and cellular PrP isoforms are encoded by the same chromosomal gene. Cell. 1986;46:417–28.

5. Prusiner SB. Novel proteinaceous infectious particles cause scrapie. Science. 1982;216:136–44.

6. Bartz JC. Prion Strain Diversity. Cold Spring Harb Perspect Med. 2016 Dec 1;6(12).

7. Telling GC, Scott M, Hsiao KK, Foster D, Yang SL, Torchia M, et al. Transmission of Creutzfeldt-Jakob disease from humans to transgenic mice expressing chimeric human-mouse prion protein. Proc Natl Acad Sci U S A. 1994 Oct 11;91(21):9936–40.

8. Telling GC, Scott M, Mastrianni J, Gabizon R, Torchia M, Cohen FE, et al. Prion propagation in mice expressing human and chimeric PrP transgenes implicates the interaction of cellular PrP with another protein. Cell. 1995 Oct 6;83(1):79–90.

9. Will RG, Ironside JW, Zeidler M, Cousens SN, Estibeiro K, Alperovitch A, et al. A new variant of Creutzfeldt-Jakob disease in the UK. Lancet. 1996;347:921–5.

10. Telling GC, Parchi P, DeArmond SJ, Cortelli P, Montagna P, Gabizon R, et al. Evidence for the conformation of the pathologic isoform of the prion protein enciphering and propagating prion diversity. Science. 1996;274:2079–82.

11. Bessen RA, Marsh RF. Distinct PrP properties suggest the molecular basis of strain variation in transmissible mink encephalopathy. J Virol. 1994;68:7859–68.

12. Benestad SL, Telling GC. Chronic wasting disease: an evolving prion disease of cervids. Handb Clin Neurol. 2018;153:135–51.

13. Williams ES, Young S. Chronic wasting disease of captive mule deer: a spongiform encephalopathy. J Wildl Dis. 1980;16:89–98.

14. Sohn HJ, Kim JH, Choi KS, Nah JJ, Joo YS, Jean YH, et al. A case of chronic wasting disease in an elk imported to Korea from Canada. J Vet Med Sci. 2002 Sep;64(9):855–8.

15. Kim TY, Shon HJ, Joo YS, Mun UK, Kang KS, Lee YS. Additional cases of Chronic Wasting Disease in imported deer in Korea. J Vet Med Sci. 2005 Aug;67(8):753–9.

16. Lee YH, Sohn HJ, Kim MJ, Kim HJ, Park KJ, Lee WY, et al. Experimental chronic wasting disease in wild type VM mice. J Vet Med Sci. 2013;75(8):1107–10.

17. Benestad SL, Mitchell G, Simmons M, Ytrehus B, Vikoren T. First case of chronic wasting disease in Europe in a Norwegian free-ranging reindeer. Vet Res. 2016 Sep 15;47(1):88.

18. Osterholm MT, Anderson CJ, Zabel MD, Scheftel JM, Moore KA, Appleby BS. Chronic Wasting Disease in Cervids: Implications for Prion Transmission to Humans and Other Animal Species. MBio. 2019 Jul 23;10(4).

19. Bian J, Christiansen JR, Moreno JA, Kane SJ, Khaychuk V, Gallegos J, et al. Primary structural differences at residue 226 of deer and elk PrP dictate selection of distinct CWD prion strains in gene-targeted mice. Proc Natl Acad Sci U S A. 2019 Jun 18;116(25):12478–87.

20. Bian J, Kim S, Kane SJ, Crowell J, Sun JL, Christiansen J, et al. Adaptive selection of a prion strain conformer corresponding to established North American CWD during propagation of novel emergent Norwegian strains in mice expressing elk or deer prion protein. PLoS Pathog. 2021 Jul;17(7):e1009748.

21. Browning SR, Mason GL, Seward T, Green M, Eliason GA, Mathiason C, et al. Transmission of prions from mule deer and elk with chronic wasting disease to transgenic mice expressing cervid PrP. Journal of virology. 2004 Dec;78(23):13345–50.

22. Angers RC, Seward TS, Napier D, Green M, Hoover E, Spraker T, et al. Chronic wasting disease prions in elk antler velvet. Emerging Infectious Diseases. 2009 May;15(5):696–703.

23. Kang HE, Weng CC, Saijo E, Saylor V, Bian J, Kim S, et al. Characterization of Conformation-Dependent Prion Protein Epitopes. The Journal of biological chemistry. 2012 Sep 4.

24. Peretz D, Williamson RA, Legname G, Matsunaga Y, Vergara J, Burton DR, et al. A change in the conformation of prions accompanies the emergence of a new prion strain. Neuron. 2002 Jun 13;34(6):921–32.

25. Taraboulos A, Jendroska K, Serban D, Yang S-L, DeArmond SJ, Prusiner SB. Regional mapping of prion proteins in brains. Proc Natl Acad Sci USA. 1992;89:7620–4.

26. Angers R, Christiansen J, Nalls AV, Kang HE, Hunter N, Hoover E, et al. Structural effects of PrP polymorphisms on intra-and interspecies prion transmission. Proc Natl Acad Sci U S A. 2014 Jul 29;111(30):11169–74.

27. Nonno R, Di Bari MA, Pirisinu L, D’Agostino C, Vanni I, Chiappini B, et al. Studies in bank voles reveal strain differences between chronic wasting disease prions from Norway and North America. Proc Natl Acad Sci U S A. 2020 Nov 23.

28. Davenport KA, Christiansen JR, Bian J, Young M, Gallegos J, Kim S, et al. Comparative analysis of prions in nervous and lymphoid tissues of chronic wasting disease-infected cervids. The Journal of general virology. 2018 May;99(5):753–8.

29. Prusiner SB, Scott M, Foster D, Pan K-M, Groth D, Mirenda C, et al. Transgenetic studies implicate interactions between homologous PrP isoforms in scrapie prion replication. Cell. 1990;63:673–86.

30. Scott M, Groth D, Foster D, Torchia M, Yang S-L, DeArmond SJ, et al. Propagation of prions with artificial properties in transgenic mice expressing chimeric PrP genes. Cell. 1993;73:979–88.

31. Bian J, Khaychuk V, Angers RC, Fernandez-Borges N, Vidal E, Meyerett-Reid C, et al. Prion replication without host adaptation during interspecies transmissions. Proc Natl Acad Sci U S A. 2017 Jan 31;114(5):1141–6.

